# A T1R-independent mechanism for responses to hyperosmotic sugars involves a carbonic anhydrase-sensitive mechanism in Type III receptor cells

**DOI:** 10.1101/2022.09.16.508275

**Authors:** B. Kalyanasundar, Ginger D. Blonde, Alan C. Spector, Susan P. Travers

## Abstract

Recent findings from our laboratory demonstrated that the rostral nucleus of solitary tract (rNST) retains some responsiveness to glutamate (MSG+amiloride-MSGa) and sugars in mice lacking the canonical T1R receptors for these tastants. Here, we recorded from the parabrachial nucleus (PBN) in mice lacking the T1R1+T1R3 heterodimer (KO1+3), using warm stimuli to optimize sugar responses and employing extended concentrations and pharmacological agents to probe mechanisms. MSGa+IMP responses were not synergized in KO1+3 mice but responses to MSGa were similar to those in B6 (WT) mice. Glutamate responses in the neurons tested were unaffected by topical application of an mGluR4 antagonist. PBN T1R-independent sugar responses, including those to concentrated glucose, were more evident than in rNST. Sugar responses were undiminished by phlorizin, an inhibitor of SGLT, a component of a hypothesized alternative glucose-sensing mechanism. There were no sugar/umami “best” neurons in KO1+3 mice, and instead, sugars activated cells that displayed acid and amiloride-insensitive NaCl responses. In WTs, concentrated sugars activated “sugar/umami” cells but also electrolyte-sensitive neurons. The efficacy of hyperosmotic sugars for driving neurons broadly responsive to electrolytes implied an origin from Type III taste bud cells. To test this, we used the carbonic anhydrase (CA) inhibitor dorzolamide (DRZ), previously shown to inhibit amiloride-insensitive sodium responses arising from Type III cells. Dorzolamide had no effect on sugar-elicited responses in WT sugar/umami PBN neurons but strongly suppressed them in WT and KO electrolyte-generalist neurons. These findings suggest a novel T1R-independent mechanism for hyperosmotic sugars, involving a CA-dependent mechanism in Type-III taste bud cells.

**Significance Statement:** Since the discovery of the *Tas1r* gene family that encodes receptors that lead to sweet and umami taste perception two decades ago, evidence has accrued that mice lacking these receptors maintain some degree of behavioral, physiological, and neural responsiveness to sugars and monosodium glutamate. But the basis for the remaining sensitivity to these nutritionally relevant compounds has remained elusive. Here we recorded from parabrachial nucleus taste neurons. Glutamate responses in mice lacking T1R1+T1R3 were unaffected by oral treatment of mGluR4 receptor antagonist suggesting that some T1R-independent glutamate responses are mediated by a different transduction pathway. Moreover, we identified a novel basis for T1R-independent responsiveness to hyperosmotic sugars that relies on carbonic anhydrase-mediated mechanism found in Type III taste bud cells.

## Introduction

Sugars and amino acids are major food components crucial for the gustatory system to detect, thereby promoting optimal nutrition. There is much evidence for a key role of the Type-I taste receptor (T1R) heterodimers, T1R2+T1R3 for sugars and T1R1+T1R3 for amino acids, in the oral detection of these molecules, since genetic deletion of one or both components of these heterodimers greatly reduces behavioral and neurophysiological responses to these stimuli. However, there is also evidence for residual sensitivity (Damak et al., 2003; Zhao et al., 2003; Delay et al., 2006; Treesukosol et al., 2009; Zukerman et al., 2009; Treesukosol et al., 2011b; Treesukosol and Spector, 2012; Smith and Spector, 2014; Glendinning et al., 2015; Sukumaran et al., 2016; Glendinning et al., 2017; Blonde et al., 2018; Glendinning et al., 2020).

We recently recorded from the first central gustatory relay, the rostral nucleus of the solitary tract (rNST), in double-knockout (double-KO) mice lacking T1R heterodimers (Kalyanasundar et al., 2020). In the absence of T1R2+T1R3 or T1R1+T1R3, neurons preferentially responsive to sugars and umami (MSG+amiloride [MSGa] + IMP) could not be found nor was the response to glutamate (MSGa) enhanced by 5’-ribonucleotides (“umami synergism”). Nevertheless, sugars and glutamate at high concentrations still activated KO neurons. Notably, in WT C57BL/6 (B6) and 129/SvJ (S129) controls, sucrose (300 mM) elicited responses comparable to those elicited by monosaccharides (1000 mM), whereas in the KO mice the monosaccharides were more effective. These data were interesting in light of previous suggestions of a T1R-independent glucose-specific mechanism functionally important in cephalic phase insulin release and in discriminating this metabolically advantageous sugar from other monosaccharides (Grill et al., 1984; Glendinning et al., 2015; Schier and Spector, 2016; Glendinning et al., 2017; Schier et al., 2019; Glendinning et al., 2020; Yasumatsu et al., 2020). However, at odds with a selective mechanism, our observations suggested equivalent T1R-independent responsiveness to glucose and fructose. Moreover, residual responses to both monosaccharides primarily occurred as sideband responses in neurons more robustly activated by electrolytes, providing little evidence for specificity (Kalyanasundar et al., 2020). Further exploration of these residual sugar responses was hampered by their small magnitude and because they occurred in a relatively small population of cells.

In the current study, we extended our investigation of residual sugar and amino acid responses in T1R1+T1R3 KO mice in several important ways. First, we applied multiple concentrations and recorded from the 2nd-order gustatory relay, the PBN, where gustatory responses are typically larger (Van Buskirk and Smith, 1981; Nakamura and Norgren, 1991; Di Lorenzo and Monroe, 1997; Geran and Travers, 2009). Second, we also presented tastants at 30o C, instead of room temperature, because warming often enhances the magnitude of neural responses and the perceived intensity of sweet stimuli (Lemon, 2017, 2021). Third, because previous studies suggest the possibility that the mGluR4 receptor contributes to glutamate taste (Chaudhari and Roper, 1998; Chaudhari et al., 2000) and that the sodium-glucose transporter, SGLT1, can serve to detect glucose-containing sugars (Yasumatsu et al., 2020), we used antagonists to these receptors to test whether they affected PBN glutamate and sugar responses.

Similar to the NST, in the PBN there were salient responses to glutamate in the T1R1+T1R3 KO mouse. However, these were unaffected by the mGluR4 antagonist, (RS)-α-methylserine-O-phosphate (MSOP). In contrast to the NST, there was a sizeable population of PBN neurons with robust T1R-independent sugar responses, but as in NST, these responses occurred primarily in electrolyte generalist cells. At equimolar, hyperosmotic concentrations, neural activity elicited by monosaccharides and disaccharides in the KOs was comparable, providing little evidence for sugar selectivity. Moreover, T1R-independent sugar responses were unaffected by the SGLT antagonist, phlorizin, but surprisingly, were dramatically suppressed by the carbonic anhydrase (CA) inhibitor, dorzolamide (DRZ). These results suggest a previously unsuspected basis for T1R-independent responses to hyperosmotic sugars, involving a CA-dependent mechanism arising from Type-III taste bud cells.

## Methods

### Animals

Our previous observations in the rNST showed that the effects on sugar and amino acid responses were virtually identical in T1R1 + T1T3 and T1R2 + T1R3 double-knockout mice, confirming the T1R3 protein as a critical heterodimeric component for neural responsiveness to both amino acids and sugars. Thus, in the present study, to be efficient, we focused on T1R1 + T1R3 KO mice (hereafter called “KO1+3”).

Subjects were adult male (N=27) and female (N=41) KO1+3 mice and B6 (C57BL/J6-Jackson Laboratory, Bar Harbor, ME) mice as WT controls (male: n=35; female = 38). Sex was evenly distributed across strains (χ2 = 0.97; *p =* 0.32) but due to limited availability, KO1+3 mice were older than B6 mice (329 ±90 days versus 288 ±103 days; p < 0.0001). Nevertheless, when age was included as a covariate in an ANOVA analysis that compared responses to the standard taste qualities, this variable was not significant (effect of strain: *p =* 0.77; age: *p =* 0.96; and stimuli X age: *p =* 0.17).

The generation of the double-knockout mice was described previously (Blonde et al., 2018; Kalyanasundar et al., 2020). Briefly, homozygous mice null for the *Tas1r1* or *Tas1r3* derived from S129 backcrossed with B6 mice (from Dr. Charles Zuker while at the University of California, San Diego [now at Columbia University]), were backcrossed an additional time. Subsequently, they were paired to generate mice heterozygous for both *Tas1r1* and *Tas1r3*. These heterozygous mice were used to create heterozygous, homozygous null, and wild type for *Tas1r1* and/or *Tas1r3*. These mice were then paired to generate mice homozygous null for both *Tas1r1* and *Tas1r3*. The double-KO mice were bred to generate the experimental double-KO mice (KO1+3). Analysis of known polymorphisms between B6 and S129 strains (Jackson Laboratory) concluded that the double-KO mice have the majority of their genome (70-80%) from the B6 strain. Moreover, in the rNST, using a similar array of stimuli, we found few differences between B6 and 129 mice (Kalyanasundar et al., 2020). Thus, in the present study, we used B6 mice as WT controls. Genotype was confirmed by PCR amplification of Tas1r1 and Tas1r3 for all double-KO animals and a few random WT mice (Transnetyx, Inc., Cordova, TN) after the experiment. The animals were maintained in a temperature and humidity-controlled colony room on a 12-h: 12-h light/dark cycle and were given ad libitum access to food and water. The Institutional Laboratory Animal Care and Use Committee at the Ohio State University approved all experimental procedures.

### Surgery

Animals were deeply anesthetized with urethane (1 g/kg, I.P.) and maintained at this level using supplemental doses of 0.25 g/kg. Isoflurane was supplemented at 2% for the initial surgery induction and continued at 0.5% until recording. Body temperature was kept at ~37°C using a heating pad and a tele-thermometer. The trachea was dissected posterior to the larynx for tracheotomy, and a length of polyethylene tubing (PE60) was inserted to aid respiration. The hypoglossal nerves were transected bilaterally to limit oromotor reflexes during testing. Four sutures were secured on the corners of the lips, and a suture was passed superficially on the tongue anterior to the foliate papillae to examine and facilitate the stimulus flow (Norgren et al., 1989; Frank, 1991; Travers and Norgren, 1995; Breza and Travers, 2016; Kalyanasundar et al., 2020). After tracheotomy, each mouse was fixed in a stereotaxic apparatus equipped with a non-traumatic head holder (custom-modified Kopf). An incision was made (~1cm) to expose the scalp. The skull was leveled in the sagittal and coronal planes by adjusting the knob of the head holder and bite piece. Then a hole was drilled (~5 mm) on the intraparietal bone to expose inferior colliculus (IC) and cerebellum (CB).

### Data Collection

Coordinates for recording from the PBN were 0.9-1.4 mm caudal and 1.0-1.4 mm lateral to lambda. A tungsten microelectrode (1.0–3.0 MΩ, FHC, ME, USA) was driven through the inferior colliculus (IC), at a right angle to the skull surface (Tokita et al., 2012) using a hydraulic microdrive (model no. 640; David Kopf Instruments) while observing multiunit neural responses with an oscilloscope and audio monitor. Neuronal activity was amplified and filtered (10,000X, 0.6–3 kHz), sampled at 10 kHz, and monitored using the SPIKE 2-CED data acquisition system (model no. 140; Cambridge Electronic Design, Cambridge, UK). Taste-responsive activity was usually encountered at 2.7-3.2 mm ventral to the surface of IC. To search for single gustatory-responsive neurons, we stimulated the whole mouth with a taste mixture (0.3 M sucrose, 0.1 M NaCl, 0.01 M citric acid, and 0.01 mM cycloheximide), artificial saliva (AS), and various individual stimuli. After isolating a single unit in the PBN, a battery of taste solutions, applied to the whole oral cavity, was tested (See Tastant and Stimulation for details) using a pressurized flow system controlled by CED software. After initial testing, the receptive field of the neuron was assessed by stroking specific taste bud subpopulations with a precision applicator brush (S379, Parkell Inc.) soaked with taste mixture. The receptive field was defined as anterior [A-fungiform papillae (FUN) and nasoincisor ducts (NID)], posterior-[P-soft palate (SP), foliate papillae (FOL), and circumvallate papillae (CV)] or mixed [M-both anterior (A) and posterior (P)) (Geran and Travers, 2006, 2009; Geran and Travers, 2013; Kalyanasundar et al., 2020). Afterwards, to mark the recording location, we made a small electrolytic lesion at the dorsal border of PBN and/or ventrally to the nucleus, or in a minority of instances, at the recording site. At the conclusion of the experiment, each animal was injected with a lethal dose of anesthesia (80 mg/kg ketamine and 100 mg/kg xylazine) and intracardially perfused using phosphate-buffered saline (PBS) and 4% paraformaldehyde (in 0.1 M phosphate buffer) containing 1.4% L-lysine acetate and 0.2% sodium meta periodate (McLean and Nakane, 1974). Subsequently, the brain was extracted, fixed overnight in 20% sucrose paraformaldehyde, blocked in the coronal plane, and stored in a sucrose-phosphate buffer before sectioning. Two series of 40 μm coronal PBN sections were cut using a freezing microtome and stored at −20°C in cryoprotectant until further use.

### Tastants

Taste solutions were made from reagent-grade chemicals (obtained from Sigma unless otherwise mentioned) dissolved in artificial saliva (AS; (Breza et al., 2010; Kalyanasundar et al., 2020). Concentrations were selected from our previous rNST study in T1R double-KO mice (Kalyanasundar et al., 2020). The experiment proceeded in several stages using different, but overlapping, sets of stimuli (summarized in **Table 1**). The stimuli common to each stage, used to test all neurons, are referred to as the “core” stimuli: these were prototypical representatives of the four classic taste qualities (“sweet”, sucrose 300 mM and glucose 1000 mM, “salty”, NaCl 100 mM, “sour”, citric acid 10 mM and “bitter”, a quinine 2.7 mM + cycloheximide 0.01 mM cocktail). In most cases, we also used 100 mM NaCl plus 100 μM amiloride (N100+a) to facilitate classifying salt-sensitive neurons into amiloride-sensitive and amiloride-insensitive groups.

**Table 1.** Stimuli, concentrations, and the number of neurons sampled for each tastant.

|  |  |  |  |  |  |
| --- | --- | --- | --- | --- | --- |
| **STIMULUS** | **CONCENTRATIONS** | **ACRONYM** | **B6** | **KO1+3** | **Refer** |
| **Core Stimuli** |  |  | **N's** |  |  |
| Bitter mixture <sup>1</sup> | 2.7 mM Q+0.01 mM CY | **B3** | 73 | 68 | Figs. 1, 2, 4 & 5 |
| Citric acid | 10 mM | **I10** |  |  | Figs. 1, 2, 4 & 5 |
| NaCl | 100 mM | **N100** |  |  | Figs. 1, 2, 4 & 5 |
| Sucrose | 300 mM | **S300** |  |  | Figs. 1, 2, 3, 4 & 5 |
| Glucose | 1000 mM | **GLU1000** | 73 | 67 | Figs. 1, 2, 3, 4 & 5 |
| **Extension** |  |  |  |  |  |
| NaCl + amiloride | 100 mM NaCl + 100 $\mu$ M a | **N100+a** | 60 | 52 | Figs. 2, 4 & 5 |
| **Saccharides (hyperosmotic high concentrations)** |  |  |  |  |  |
| Sucrose | 600 mM | **S600** | 52 | 49 | Figs. 3 & 5 |
| Glucose |  | **GLU600** | 27 | 18 | Figs. 3 & 5 |
| Fructose |  | **FRU600** | 27 | 17 | Figs. 3 & 5 |
| Fructose | 1000 mM | **FRU1000** | 12 | 10 | Fig. 4 |
| **Artificial sweetener** |  |  |  |  |  |
| Na. Saccharin | 5 mM | **SACC5** | 10 | 10 | Fig. 4 |
| **Sodium Glucose co-transporter antagonist (SGLT I/II)** |  |  |  |  |  |
| Phlorizin | 1 mM | **Phlo** |  |  |  |
| Glucose+Phlorizin | 1000 + 1 mM | **GLU100+Phlo** | 8 | 3 | Fig. 4 |
| Fructose+Phlorizin | 1000+ 1 mM | **FRU1000+Phlo** | 8 | 3 | Fig. 4 |
| **Carbonic anhydrase inhibitor** |  |  |  |  |  |
| Dorzolamide | 0.5% | **DRZ** | 11 | 7 | Fig. 5 |
| **Glutamate & umami** |  |  |  |  |  |
| MSG<sup>2</sup>+amiloride<sup>3</sup> | 50 or 300 or 600 mM MSG + 100 $\mu$ M a | **MSGa (50/300/600)** | 59/67/53 | 49/60/50 | Fig. 6 |
| Inosine- 5'-monophosphate | 2.5 mM | **IMP2.5** | 29 | 26 | Fig. 6 |
| umami | 50 or 300 or 600 M MSG + 100 $\mu$ M a + 2.5 mM IMP | **umami (50/300/600)** | 60/69/55 | 53/66/52 | Fig. 6 |
| **Metabotropic glutamate receptor (mGLUR4) Antagonist** |  |  |  |  |  |
| (RS)-α-Methylserine-O-phosphate | 0.5 mM | **MSOP** | | | |
| umami + MSOP | 300 mM MSG + 100 $\mu$ M a + 2.5 mM IMP + 0.5 mM MSOP | **U300+MSOP** | 2 | 3 | Fig. 6 |
| MSGa + MSOP | 6000 mM MSG + 100 $\mu$ M a + 0.5 mM MSOP | **MSGa600 + MSOP** | 0 | 4 | Fig. 6 |

<sup>1</sup> bitter mixture: quinine monohydrochloride (Q) + cycloheximide (CY), <sup>2</sup> monosodium glutamate [MSG], <sup>3</sup> amiloride [a]
All solutions were prepared in the Artificial saliva (composition based on Breza et al. 2010)

Additional sugar concentrations were tested, as well as select orally applied pharmacological agents to probe potential T1R-independent transduction pathways. Thus, we tested a subset of neurons with an equimolar concentration (600 mM) of sucrose, glucose and fructose. In addition, we employed phlorizin (a competitive inhibitor for SGLT1/2, purchased from Sigma-Aldrich) using a protocol and concentration (1 mM) previously shown to affect T1R-independent sugar responses in the chorda tympani and glossopharyngeal nerves (Yasumatsu et al., 2020). After assessing sugar and other responses in the control state, the oral cavity was bathed with phlorizin (1 mM) for 5 minutes and then the stimuli were tested again mixed in a cocktail with phlorizin. In other experiments, we used a cell-permeant broad-spectrum carbonic anhydrase inhibitor Dorzolamide (DRZ). Inhibiting carbonic anhydrase with DRZ has previously been demonstrated to block a subset of amiloride-insensitive sodium responses in Type III taste bud cells (Lewandowski et al., 2016) and those in the CT nerve that probably arise from such cells (Oka et al., 2013). We used DRZ at a concentration of 0.5%, the same as used topically by (Oka et al., 2013) to study CT responses. Although this manipulation has not previously been shown to interfere with sugar responses, we postulated that this was a possibility based on our observations that sugar responses in the PBN in the KO1+3 mice occurred in neurons that displayed broad electrolyte sensitivity, including amiloride-insensitive sodium responses and thus presumably received some input from Type III cells.

Finally, to gain a fuller appreciation of T1R-independent glutamate responses, we tested a concentration series (50, 300 and 600 mM) of monosodium glutamate (MSG) + 100 μM amiloride (a) + 2.5 mM inosine 5’-monophosphate (IMP)] and MSG+a (MSGa). Hereafter, for convenience, we often refer to the cocktail with IMP as “umami”, the perceptual quality that this mixture reportedly generates in humans. To refer to general effects that include both the umami stimulus and MSGa, we use the term “glutamate”. The concentrations of MSGa and MSGa+IMP were selected based on our previous rNST recordings, where responses to 600 mM umami were unaffected but responses to 100 mM were strongly suppressed by T1R gene deletion. Thus, 300 mM was used to fill in the range between the affected and unaffected concentrations and 50 mM and 600 mM served to verify the ends of the range. Further, to validate the function of the T1R1+T1R3 receptor in umami synergism, we tested 2.5 mM IMP alone along with the three (50, 300, and 600 mM) concentrations of MSGa with and without IMP. To understand the role of metabotropic glutamate receptor (mGluR) for the T1R-independent glutamate responsiveness, in selected cases where umami or MSGa responses were prominent, we used a cocktail of the mGluR-4 antagonist [0.5 mM MSOP: (RS)-α-methylserine-O-phosphate obtained from Tocris Biosciences] with either umami at 300 mM or MSGa at 600 mM. The choice of MSOP as an antagonist and its concentration was based on previous studies demonstrating marked inhibition of mGluR agonist [L-(+)-2-amino-4-phosphonobutyrate: L-AP4] mediated responses of taste bud cells of T1R3-KO mice (Pal Choudhuri et al., 2016).

### Stimulation

Two major modifications were made in the stimulation protocol in these PBN recordings compared to our previous rNST experiments in KO1+3 mice (Kalyanasundar et al., 2020). All taste stimuli, including AS, were presented at near-physiological temperature (30oC), instead of room temperature, in our attempt to augment residual sugar responses in the double-KO mice. Also, to maintain a constant temperature, we continuously delivered AS between taste stimulations, instead of incorporating a pause in fluid flow. Stimuli were kept in air-tight, pressurized glass bottles connected to separate tubes, each regulated by a computer-controlled solenoid valve. Tubes from the different stimuli met at a 16:1 manifold (Warner Systems) near the mouth and then passed through an in-line heater (SH-27B; Warner Instruments, CT, USA) which exited to a “L-Shaped” glass capillary (1.2 mm inner diameter and length ~10 cm) positioned into the mouse oral cavity. The in-line heater included a thermistor that provided feedback to the temperature controller (TC-344C, Warner instruments, CT, USA). A second thermistor probe was placed adjacent to the end of the glass capillary where the stimulus entered the mouth to monitor the actual temperature of the delivered tastants. Due to the presence of the in-line heater, there was a 1-s (mean ± SD: 1.08 ± 0.08s) lag after valve opening before the stimulus reached the mouth. For simplicity, however, we used the valve opening and closing times for analysis.

Once a single unit was isolated, testing was initiated by adapting the oral cavity to a continuous flow (0.3 ml/s) of ~30o AS until a stable temperature was attained. Subsequently, we presented stimuli in a varied order, at a rate of 0.3 ml/s with the stimulator positioned so that fluids bathed the entire oral cavity, as monitored through an operating microscope. Each stimulus was presented for 10-s. The interstimulus interval, during which AS flowed, was at least 60-s. However, sometimes we waited for another minute or so (~120-s total) between stimuli to recover baseline spontaneous activity, which was often reduced when the epithelial sodium channel (ENaC) blocker (amiloride), was included in the stimulus. Stimulations were repeated whenever possible. After testing all the stimuli, the receptive field was evaluated as described above.

### Data analysis

Spikes were sorted and counted for each trial using Spike 2 software (CED Cambridge, Ltd.). Statistical analysis and quantification were performed using Excel (Versions 2013 and 2016), Systat (Version 13.1), and Graph pad Prism 9 (version 9.3.0). The mean spontaneous and 10-s AS pre-rinse response rates across trials were obtained for each neuron. The response measure was the number of spikes in the 10-s stimulation period minus the preceding 10-s AS (net spikes/10-s). A significant response was defined as one where the response was ≥ 1 spike/s and ≥ 2.5 times SD of the mean response to the 10-s AS pre-rinse (Nishijo et al., 1991; Geran and Travers, 2009; Kalyanasundar et al., 2020). Averages were used when taste stimulation was repeated. The proportions of significant responses for each stimulus were determined across strains and compared using χ2 tests. Differences in response magnitude were assessed using one- or two-way mixed ANOVAs with post-hoc testing accomplished with Tukey’s or Bonferroni-adjusted paired T-tests. Significance was set at *p* ≤ 0.05 for all comparisons. Errors and error bars are means ± SE. To facilitate the clarity of the text, statistical details are usually presented in the Figure captions.

Neurons were divided into chemosensitive groups with hierarchical cluster analysis (Pearson’s correlations and an average amalgamation schedule) using the core stimuli (see Table 1 for details) and NaCl to which the (ENaC) blocker, amiloride was added (N100+a) to differentiate amiloride-sensitive and insensitive groups of cells. The breadth of tuning across the core stimuli was quantified using the noise: signal ratio (Spector and Travers, 2005) and the entropy measure (Smith and Travers, 1979). For neurons where IMP was tested alone (B6: n=29 and KO1+3: n=26), we calculated the synergistic ratio as the magnitude of the response to umami divided by the sum of responses to MSGa and IMP alone. The criterion for synergism was set at 1.2 as previously defined (Ninomiya and Funakoshi, 1989; Tokita and Boughter, 2012; Kalyanasundar et al., 2020).

### Brief-access taste test

To evaluate the relative efficacy the sugars and concentrations used for neurophysiological testing to motivate licking, brief-access taste tests were conducted in a chamber that allowed detection of licking during presentation of multiple stimuli in discrete trials (known as the “Davis Rig”; Davis MS160-Mouse; DiLog Instruments, Tallahassee, FL). Water-deprived (~21-24 h) mice were initially (days 1 and 2) trained to lick for water in daily 30-min sessions at a single spout in the rig. To accustom the mice to the trial structure, on days 3 and 4, we presented water from multiple sipper tubes during sequential 10-s trials with intertrial intervals of 7.5-s. After each session, each mouse was rehydrated in its home cage for 20 minutes with ad libitum access to food, followed by overnight water deprivation before the next session. During the actual taste tests, mice were water- and food-restricted by providing a daily ration of 2 ml of water and 1 g of chow in fresh bedding. This regimen maintained them at ~85% of their ad libitum body weight (Glendinning et al., 2002). To match the neurophysiological study, a total of 5 different sugar solutions: 300 mM sucrose, 600 mM sucrose, fructose, and glucose, and 1000 mM glucose, along with water were presented to naïve B6 (N=5, M-3, and F-2) mice. This array of sugars was presented in randomized blocks of 10-s trials across the 30-min session; the test session was repeated once. Lick scores (licks to stimulus – mean session licks to water) were calculated for each stimulus and the means derived across sessions for a given animal. ANOVA followed by post-hoc pairwise comparisons (t-tests) were made across sugars.

### Histology and immunohistochemistry

To define recording sites relative to the cytoarchitecture of the PBN, we stained the first series of sections with Cresyl violet (Deitch and Moses, 1957). We immunostained a second series with Satb2, which is expressed by many neurons in the taste-responsive “waist area”, or double-labeled for NeuN or Satb2 in combination for cGRP because this neuropeptide preferentially marks neurons in the external lateral subnuclei (Campos et al., 2018; Fu et al., 2019; Jarvie et al., 2021). Standard DAB immunohistochemistry was carried out on free-floating slices at RT unless otherwise mentioned. Sections were washed in PBS before and after treatment with 1% sodium borohydride and 0.5% H_2_O_2_. Non-specific binding sites were blocked and membranes permeabilized in a mixture of 0.3% Triton, 3% bovine serum albumin, and 7.5% donkey serum for 90-min prior to adding the primary antibod(ies) (Satb2: 1:3K ABCAM Cat#51502; RRID: AB_882455; cGRP: 1:30K, ABCAM cat#36001; RRID: AB_725807, NeuN: 1:1K, Millipore MAB377, RRID:AB_2313673).

Sections were left overnight on a shaker table at RT or at 4°C for 48 h with intermittent shaking at RT. Subsequently, sections were soaked in secondary antibody (1:500 biotinylated donkey anti-goat IgG and/or anti-mouse IgG: Jackson, Inc., Cat#705-065-147; RRID: AB_2340397 and 715-065-151; RRID: AB_2340785) diluted in blocking solution (90-min) and then in an avidin-biotin mixture (Elite Kit, Vector, Cat#PK-6100; RRID: AB_2336819) diluted in 0.1 M PB and 0.1% BSA. The chromagen reaction commenced with a 15-min incubation in 0.05% 3, 3’-diaminobenzidine-HCl (DAB) without or with 0.015% nickel ammonium sulfate, before adding H_2_O_2_ to achieve a concentration of 0.075% to produce the final reaction product. Sections were inspected under brightfield and darkfield illumination using a light microscope (Nikon E600), and digital photomicrographs taken of sections containing lesions and/or electrode tracks (Nikon DXM 1200, Act II software). Lesion locations were plotted on a representative series of Sat2b-stained sections that we prepared. Subnuclei were identified based on previous descriptions (Tokita and Boughter, 2016). Only neurons where lesions were made at the site of recording or ≤ 200μM dorsal or ventral to the recording site were reconstructed.

## RESULTS

### T1-R independent sugar responses are salient in the second-order taste relay

**Figure 1A** depicts the experimental setup. To provide a sensitive assay, we delivered warm stimuli to the entire oral cavity while recording from the taste region of the PBN. All 141 taste-PBN neurons (B6: N=73, and KO1+3: N=68) were tested with the five core stimuli which included midrange concentrations of the four standard taste qualities and 1000 mM glucose. Mean and individual responses (net spikes/10s) appear in **Figure 1B**. As expected, responses to sucrose and glucose were significantly smaller in mice lacking the T1R1+T1R3 heterodimer. In contrast, mean responses to the other qualities were larger in the KOs, although the difference only reached significance for NaCl (*p* = 0.007), and the increment for citric acid was marginal. We interpret these increases as arising from the absence of saccharide-selective neurons in the KOs. That is, average responses to the electrolytes and bitter stimuli are not “diluted” by the presence of the sugar cells, which were poorly responsive to these compounds (see below). The mean spontaneous activity of the taste-responsive neurons was comparable across strains (B6: 13.8± 2.4; and KO1+3: 19.4 ± 3.5 spikes/10-s; *p =* 0.18).

**Figure 1.**
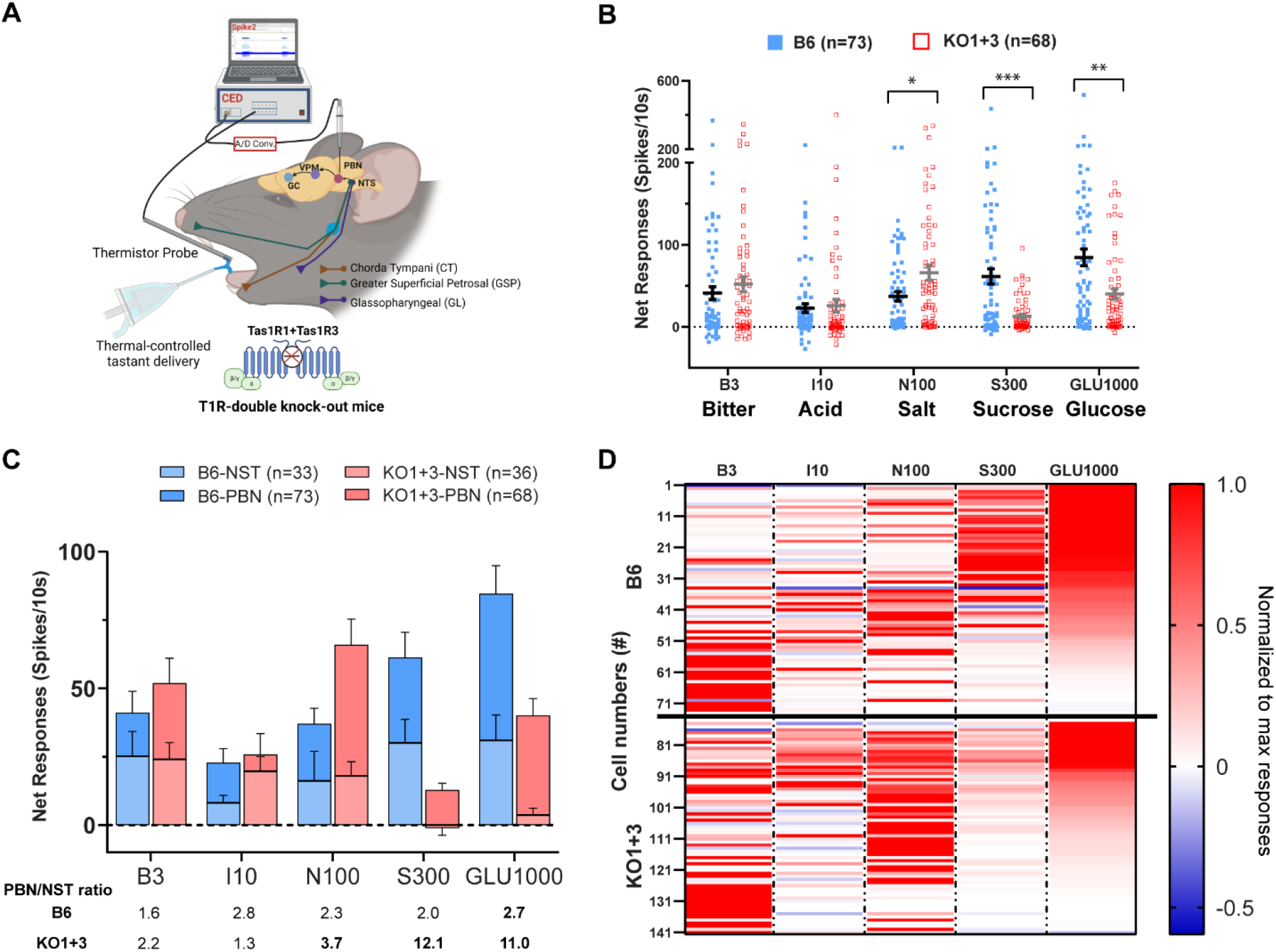
T1R-independent sugar responses were evident in the parabrachial nucleus (PBN), especially for 1000 mM Glucose. **A**. Drawing of the experimental setup for single-unit recordings from the PBN in T1R double-KO and B6 mice while stimulating the whole oral cavity with warm tastants (30°C). The three primary afferent nerves that supply lingual and palatal oral taste buds and their central projections are also shown. **B**. Net taste responses (spikes/10s) of all sampled PBN neurons from B6 (blue filled squares) and KO1+3 mice (red open squares) to the core stimuli that include four classic taste qualities (bitter: cycloheximide + quinine-B3, sour: citric acid-I10, salty: NaCl-N100, sweet: sucrose-S300 and glucose-GLU1000). Horizontal bars depict means (±SEM). An ANOVA revealed an effect of strain that approached significance (*p=*0.08), a significant effect of stimulus and a strain X stimulus interaction (both *p’s* <0.0001). Note there is a break in the y-axis. \**p* = 4.50E-02, \*\**p* = 1.48E-03 and *** 1.14E-05 between B6 and KO1+3 mice (Bonferroni-adjusted t-tests). **C**. Superimposed bar graphs comparing neural taste responses to the core stimuli between NST (data from our previous study) and PBN across B6 (NST-light blue, PBN-dark blue) and KO1+3 (NST-light red, PBN-dark red) mice. Numbers below the graph indicated the ratio of the mean response in the PBN: NST. Interestingly, the responses to sugars in the KO1+3 mice are ~10X times larger in the PBN. Bolded numbers indicate a significant difference in the response in the PBN versus the NST. (Bonferroni-adjusted t-tests). **D**. The heat map shows response profiles of all PBN neurons to the core stimuli recorded from B6 (top) and KO1+3 (bottom) mice. Responses are normalized to the maximum response for a given neuron and ordered by glucose responsiveness. Each row represents a neuron. The horizontal solid line and vertical black dotted lines separate the strain and stimuli. Note that, in the KO1+3 mice, responses to 1000 mM glucose (GLU1000) were more robust than to 300 mM sucrose (SUC300) in neurons that showed prominent responses to salt and citric acid.

Although the KO1+3 mice had diminished responses to sugars overall, they nonetheless displayed a notable number of robust responses to these compounds (**Figure 1B**). Thus, presumably because we recorded from the PBN, where gustatory responses typically are larger (Perrotto and Scott, 1976; Van Buskirk and Smith, 1981; Geran and Travers, 2013), and used warm stimuli, reported to enhance responses to sweet stimuli (Lemon, 2021), we had a more sensitive platform to probe T1R-independent sugar responses than in the NST. In fact, for both the B6 and KO1+3 mice, all the stimuli evoked nominally larger responses in PBN than in NST neurons although only some of these increments were statistically significant (**Fig. 1C**). It was particularly notable that 55% of PBN neurons in the KO1+3 mice responded significantly to glucose compared to 18% in the medulla and that both sugars elicited mean responses in the PBN of double-KO mice that were over 10X greater than in NST.

To gain insight into the nature of the T1R1+3-independent sugar responses, we made a heat map of the responses to the core stimuli, in which we ordered neurons according to glucose responsiveness across both strains (**Fig. 1D)**. Interestingly, in both B6 and KO1+3 mice, a sizable proportion of cells (0.23 and 0.14) responded better or comparably to glucose compared to the other stimuli. However, it seemed apparent that the chemosensitive profiles of the “glucose-best” neurons differed dramatically by strain; in the KO they exhibited prominent responses to NaCl and citric acid, whereas many in the comparable WT group had responses more restricted to sucrose. To systematically explore this, we performed a cluster analysis encompassing both WT and KO1+3 neurons using these 5 stimuli and NaCl + amiloride (N100+a) to further differentiate amiloride-sensitive versus insensitive NaCl-responsive cells.

### T1R-independent sugar responses occur in electrolyte generalist neurons

We identified five major chemosensitive neuron types (**Figure 2A**). The color-coding on the dendrogram identifies the optimal core stimulus for a given cell. Except for neurons selectively responsive to sugars, each type was evenly distributed in both strains (**Fig. 2B**; χ2 = 4.88, *p=*0.18**). Figure 2C** indicates that a given cluster exhibited a similar response profile regardless of the presence of T1R1+T1R3; ANOVA indicated a lack of a 3-way interaction between stimulus X cluster X strain (see Figure caption for details). The “sugar” group, observed only in B6 mice, was highly selective for sugar stimuli (**Fig. 2D**) and, on average, responded optimally and comparably to 300 mM sucrose and 1000 mM glucose but hardly to the tastants representing other qualitative classes. Notably, only one other group comprised neurons that, on average, responded optimally to 1000 mM glucose. Unlike the sugar group, this cluster included neurons from both B6 and KO1+3 mice. These cells responded second-best and comparably to NaCl and NaCl + amiloride (Na); i.e., they were amiloride-insensitive. Therefore, we refer to them as “EGNai”. EGNai neurons were broadly tuned across qualities **(Fig. 2D)** and resemble electrolyte generalist neurons described in many previous single-unit taste studies (Frank et al., 1983; Ninomiya and Funakoshi, 1988; Contreras and Lundy, 2000; Breza et al., 2010; Kalyanasundar et al., 2020).

**Figure 2.**
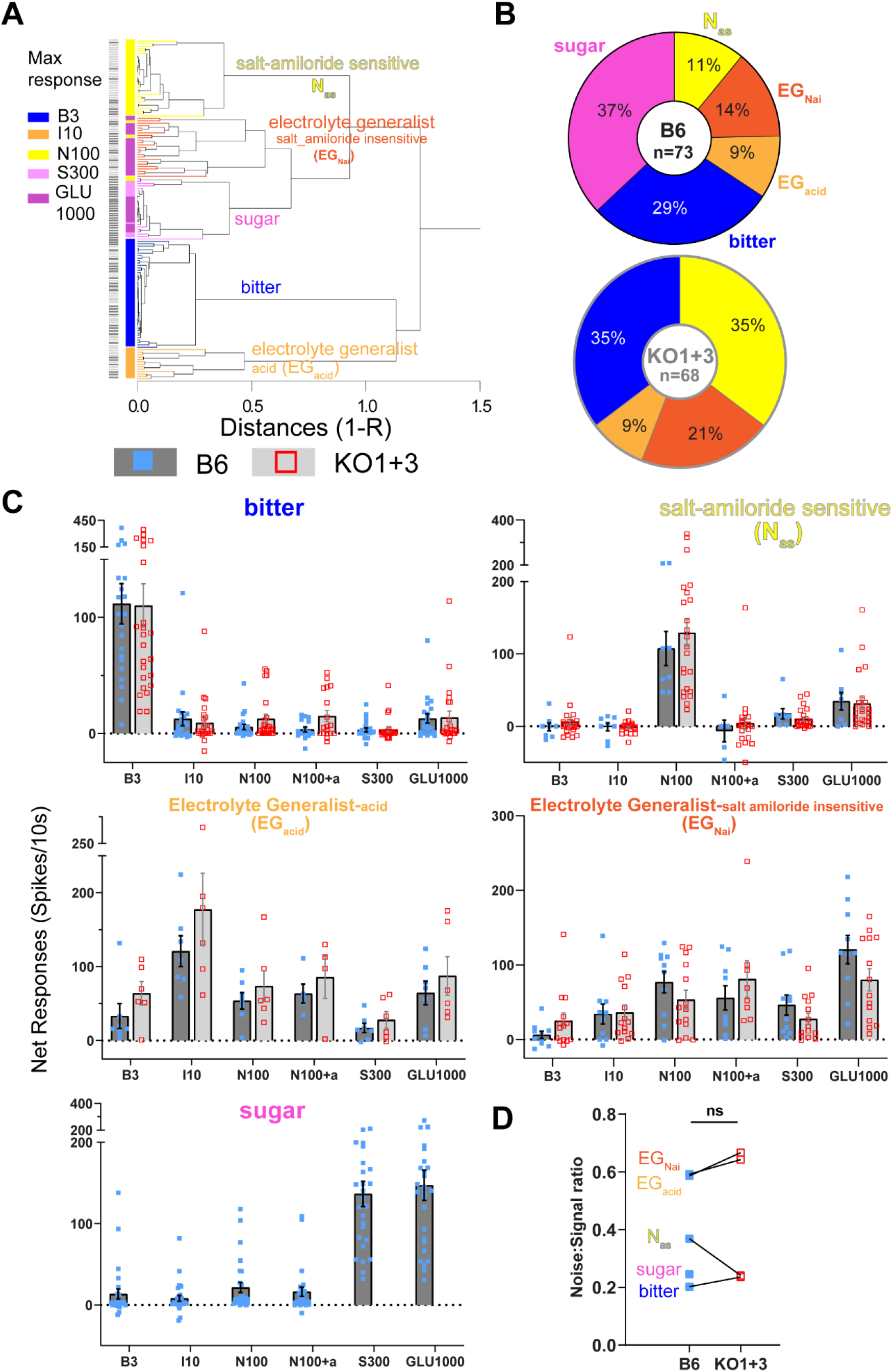
“Sugar”-best neurons were absent in the PBN of T1R double-KO mice, but T1R-independent sugar responses were robust in electrolyte generalist neurons responsive to amiloride insensitive salt and acid. **A**. Dendrogram of all PBN (n=141) neurons from both strains. Hierarchical cluster analysis (Pearson’s r, average linkage) included the core stimuli (B3, I10, N100, S300, and GLU1000) and NaCl plus amiloride (N100+a). Five chemosensitive groups were identified: bitter, salt amiloride-sensitive (N_as_), electrolyte generalist acid (EG_acid_), electrolyte generalist salt-amiloride-insensitive (EG_Nai_), and sugar. (Note that, when tested, the sugar neurons also responded well to umami stimuli; we present these data later in the manuscript). The distance metric on the x-axis is 1-r. Black (B6) and grey (KO1+3) bars on the y-axis specify mouse strains. Color bars on the y-axis indicate the best stimulus. Note that EG_Nai_ neurons responded best to 1000 mM glucose in both strains. **B**. The pie charts show the percentage of neurons in each cluster for B6 (left) and KO1+3 (right) mice. The KO1+3 mice are missing sugar cluster neurons (N=141; χ2=35.81, *p* < 0.0001), while the distribution of the other cluster types was equivalent between strains (n=114; χ2=4.88, *P=* 0.18). **C**. Mean (±SEM) net responses (spikes/10s) for the five chemosensitive neuron types across all the stimuli for B6 (dark bars) and KO1+3 (grey bars) mice. Symbols in each bar represent the individual responses to each stimulus (B6-blue squares; KO1+3 mice-red open squares). Note that y-axis scales are different for cluster types. For the clusters that were present in both strains, responses across stimuli were highly similar; i.e., there was no effect of strain nor an interaction between stimulus, cluster and strain (ANOVA: main effect of strain: *p=* 0.44, cluster: p < 0.0001, cluster X strain: *p =* 0.26, stimulus X cluster: *p* <0.0001, and stimulus X cluster X strain: *p=* 0.63). Glucose responses are prevalent in EG_Nai_ and EG_acid_ cluster neurons. **D**. The breadth of tuning across the cluster types was quantified as the noise: signal ratio (N/S) (Spector and Travers, 2005) compared across B6 and KO1+3 mice. Entropy (Smith and Travers, 1979) measure yields a similar conclusion.

Another cluster of WT and KO cells also were electrolyte generalist-like but we called them “EGacid” because, on average, they responded most robustly to citric acid. Otherwise, acid neurons were like EGNai neurons in also having robust amiloride-insensitive responses to NaCl and being broadly tuned **(Fig. 2D)**. None of the EGacid neurons responded best to glucose, but the average glucose response was 50% as large as that elicited by citric acid. Like sugar neurons, amiloride-sensitive salt (Nas) and bitter neurons were more selectively tuned to a single perceptual class of stimuli (**Figs. 2C&D**).

As mentioned above and consistent with what we observed in the rNST (Kalyanasundar et al., 2020), responses to 300 mM sucrose and 1000 mM glucose were comparable (S300 vs. GLU1000: 136.6 vs. 147.2, paired t-test: t= −1.5, *p* = 0.15) for neurons in the B6 sugar cluster **(Fig. 2C)**. Since this cluster is only present in B6 mice, it seems likely that these sugar responses are T1R-dependent. In contrast, the effectiveness of these two stimuli was different in the B6 EGNai (paired t-test: t=−6.88, *p* =0.0002) and EGacid clusters (t= −3.72, *p*= 0.018), with glucose eliciting the more prominent response, as it did in the KO1+3 mice (EGNai: t=−5.53, *p* =.0002; EGacid: t = −3.82, *p* = .024). This implies that there is a common factor driving the putative T1R1+3-independent sugar responses in PBN neurons sensitive to electrolytes in both strains of mice.

### The relative effectiveness of sucrose and glucose is different for putative T1R-dependent and -independent responses

The greater efficacy of glucose compared to sucrose in KO1+3 EG_Nai_ and EGacid neurons is consistent with the previous suggestion that glucose is a more potent stimulus for T1R-independent sugar receptors (Damak et al., 2003; Kalyanasundar et al., 2020). Alternatively, this could be due to the fact that glucose was tested at a higher concentration than sucrose. To disentangle these variables, we tested an equimolar concentration (600 mM) of sucrose, glucose, and fructose in a subset of neurons.

Figure 3. shows the individual and mean responses to this panel of 600 mM sugars, along with 300 mM sucrose and 1000 mM glucose. Responses are shown separately for B6 sugar cluster cells (**Fig. 3A;** putative T1R-dependent) and “non-sugar” B6 and KO1+3 cells (**Fig. 3C;** putative T1R-independent). For B6 sugar cells (N=15), all three sugars were differentially effective with sucrose > fructose > glucose when tested at the same concentration. However, in the non-sugar neurons (N=28, 11 WT, 15 KO cells), although fructose elicited a slightly larger response than sucrose, responses to 600 mM sucrose and glucose were comparable. This suggests that concentration is a key variable driving the larger 1000 mM glucose versus 300 Mm sucrose responses in the T1R-independent neurons. In turn, this implies that a property other than its qualitative or hedonic features may be responsible. Osmolarity may be critical, based on the observation that T1R-independent responses that achieved the response criterion were negligible at 300 mM (typical plasma osmolarity is 275-295 mOsm) but common at 600 mM. Moreover, for a given sugar (sucrose and glucose), response magnitude was a positive function of stimulus concentration in the non-sugar neurons.

To provide a behavioral context for interpreting these data, we tested all five sugars in a brief-access taste test in WT mice **(Fig. 3B)**. Notably, the order of effectiveness in promoting licking behavior matched the neural order of effectiveness for B6 sugar neurons quite well, with only one exception (300 mM sucrose was more effective than 600 mM fructose behaviorally, but the opposite was true for the neural responses) whereas the order of effectiveness in the non-sugar neurons was very different. Strikingly, at 600 mM, sucrose was the most effective stimulus for eliciting licking in B6 mice, followed by fructose and glucose, which was the same as the order of effectiveness in stimulating the putative T1R-dependent neurons, whereas these sucrose and glucose elicited comparable responses in the putative T1R-independent neurons in this strain.

**Figure 3.**
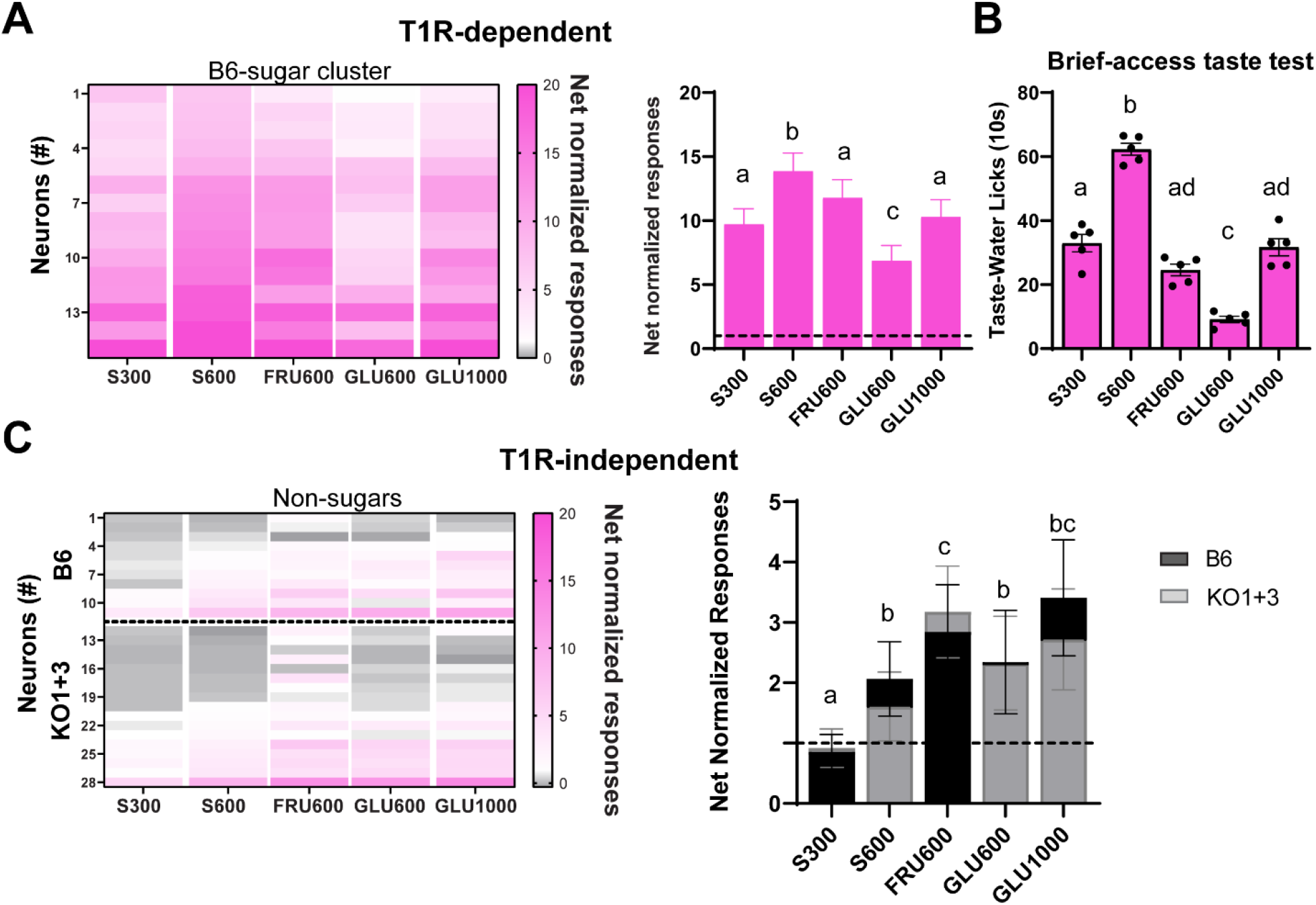
T1R-dependent responses to equimolar sucrose and glucose are markedly different in B6 sugar neurons paralleling their efficacy in the brief access taste test but are equivalent in non-sugar B6 and KO 1+3 cells. **A. Left:** Heat maps show responses for three different sugars at an equimolar concentration (600 mM) and the sugars from the core stimuli for the B6-sugar cluster neurons; sugar responses in these cells are likely to be T1R-dependent. Net responses normalized to the response criteria. **Right:** Average (±SEM) net normalized sugar responses of B6 mice sugar cluster neurons shown in left. Dotted lines represent the response criterion. **B**. Average (±SEM) lick scores (taste-water licks) for the different sugars in a brief access assay (10s trials). Only B6 mice were tested. Each dot represents an individual mouse. Notably, the order of behavioral preference mostly corresponds with T1R-dependent responses in the B6 sugar neurons. **C. Left:** Putative T1R-independent non-sugar cluster neurons in B6 (2 N_as_, 4 EG_Nai_, 2 EG_acid_, and 3 bitter neurons) and KO1+3 (7 N_as_, 5 EG_Nai_, 1 EG_acid_: 4 bitter neurons) mice. **Right:** Superimposed bar graphs (B6 [black] and KO1+3 [grey]) are the average (±SEM) net normalized sugar responses of putative T1R-independent cluster neurons. At concentrations ≥ 600 mM, sucrose, fructose and glucose all elicited mean responses that exceeded the criterion (a response that just meets criterion = 1). For both the neural and behavioral data, stimuli that differed significantly (*p* < 0.05) lack a shared letter based on Bonferroni-adjusted t-tests (data were combined for the B6 and KO1+3 non-sugar cells) following a repeated measures ANOVA.

### Blocking sodium-glucose co-transporters had no effect on T1R1+3-independent responses to sugars

A recent peripheral nerve recording study in B6 and T1R3-single KO mice suggested that the sodium-glucose co-transporter (SGLT) played a role in transducing T1R3-independent responses specific for glucose (Yasumatsu et al., 2020). Thus, in another subset of cells (single-unit: 8 B6, 4 KO1+3 and multi-unit: 2 B6 and 1 KO1+3), we used topical application of the SGLT1/2 blocker phlorizin (Phlo), at the same effective dose used by (Yasumatsu et al., 2020) to determine whether inhibiting this transporter suppressed T1R-independent glucose responses reaching the PBN. To better assess specificity for glucose, fructose and saccharin were added to the stimulus panel. **Figure 4A** shows the mean responses from the B6 and KO1+3 mice tested with phlorizin. Similar to the larger sample, sugar responses in KO1+3 mice were dramatically smaller. Moreover, saccharin responses were completely abolished in the KO1+3 mice, fructose and glucose responses were equivalent at 1000 mM without Phlo. The addition of Phlo to either fructose or glucose did not suppress taste responses in B6, or KO1+3 mice (**Fig. 4B**).

**Figure 4.**
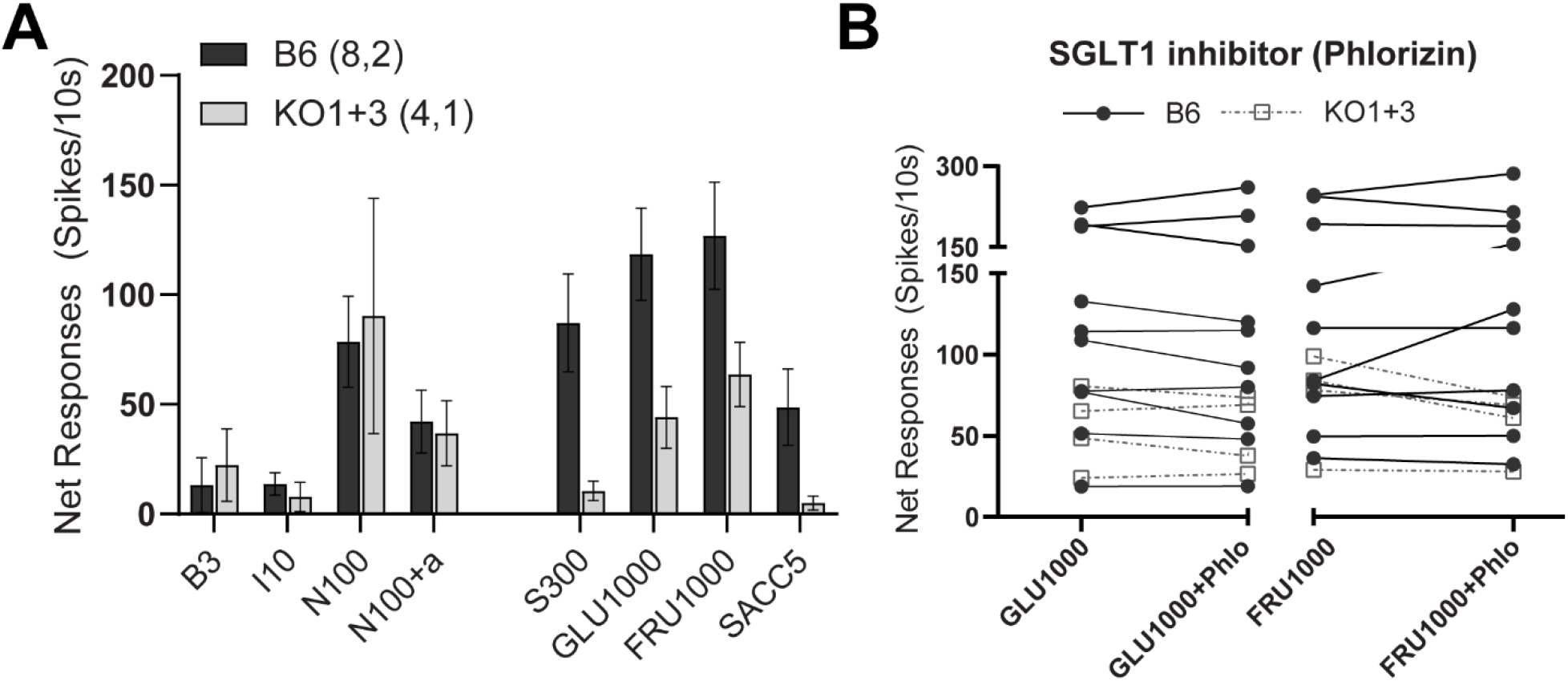
Treatment with sodium-glucose co-transporter 1 (SGLT1) antagonist did not affect responses to 1000 mM glucose or fructose. **A**. Bar graph showing average (±SEM) net responses of B6 (Black bars, N=8) and KO1+3 (grey bars, N=4) single PBN neurons (a subset those in **Figure 1**) and average multi-unit activity from three sites from the same experiments (B6: N=2 and KO1+3: N=1) tested with the core stimuli, along with 1000 mM fructose (FRU1000), and 5 mM sodium saccharin (SACC 5). All sugar responses were greatly reduced in KO1+3 mice. There was no response to saccharin in the KO mice. **B**. Connected dot plot showing the average net responses to each neuron tested with 1000 mM glucose and fructose with and without the competitive inhibitor of SGLT1, phlorizin (1 mM). Phlorizin was tested in almost all chemosensitive cluster type neurons except EG_acid_ neurons (B6-N_as_: 2, EG_Nai_: 2, bitter: 1 and sugar: 5 and KO1+3-N_as_: 2, EG_Nai_: 2 and bitter: 1). At least at these high concentrations, neither glucose nor fructose responses were affected by phlorizin.

### T1R-independent sugar responses require carbonic anhydrase activity

If the sodium-glucose transporter does not impact the T1R-independent sugar responses we observed in the PBN, additional mechanisms must contribute. Surprisingly, the chemosensitive profiles of the PBN neurons that responded to sugars in a T1R-independent manner suggested the possibility that sugars may activate a receptor mechanism and/or taste bud cell type that also transduces electrolytes, and that these signals are conveyed to broadly-tuned EG_Nai_ and EGacid PBN neurons.

Multiple lines of evidence have shown that type III taste bud cells exhibit amiloride-insensitive salt responses and are activated by acids. Moreover, previous work has demonstrated that amiloride-insensitive NaCl responses in Type III taste bud cells (Lewandowski et al., 2016) and the chorda tympani (that likely arise from Type III cells, (Oka et al., 2013) are blocked by a broad-spectrum carbonic anhydrase inhibitor, dorzolamide (DRZ). To assess whether the T1R-independent sugar responses might share this mechanism, we evaluated effects of topical incubation with DRZ (0.5%). **Figure 5A** compares responses of a KO1+3 EG_Nai_ neuron to several stimuli before and after DRZ treatment. Before treatment, this broadly tuned, amiloride-insensitive cell was most responsive to concentrated sugars (S600, FRU600, GLU600, and GLU1000) but was also activated by other stimuli. A 5 minutes lingual incubation with DRZ drastically suppressed responses to all the sugars as well as to NaCl (with or without amiloride, **Fig. 5A**, right panel). However, acid and bitter-driven activity was unaffected.

**Figure 5.**
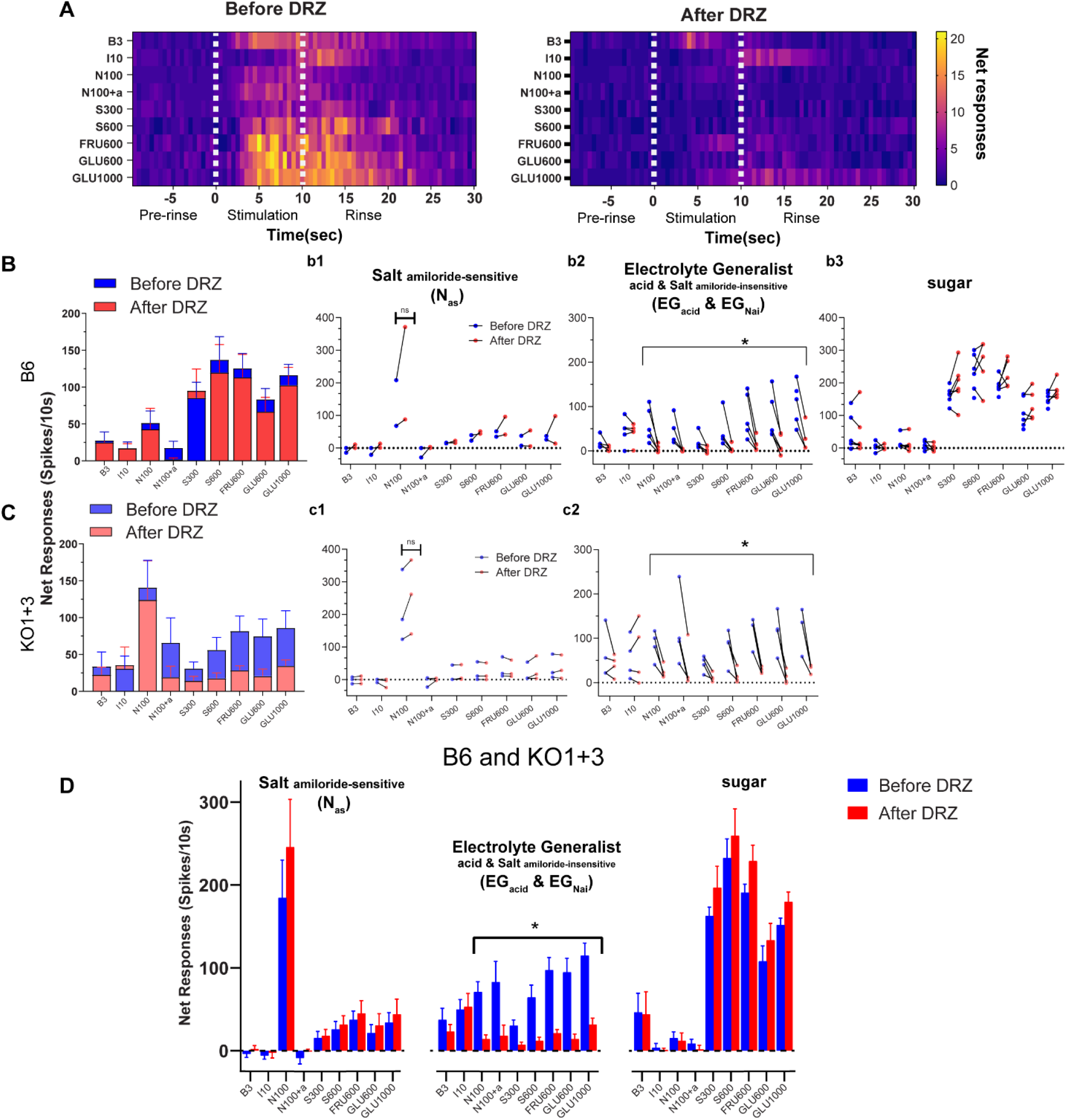
A carbonic anhydrase inhibitor, dorzolamide suppresses T1R-independent sugar responses. **A**. Heat map showing the time course (500ms bins) of responses from a KO1+3 neuron to various stimuli before (left) and 5 minutes after (right) the oral application of dorzolamide (DRZ-0.5% dissolved in artificial saliva). Vertical white dotted lines indicate the taste stimulation period (10-s). This neuron was classified as an EG_Nai_ neuron, DRZ treatment suppressed the T1R-independent responses to all sugars along with NaCl and NaCl + amiloride responses. Note that the response to acid (I3) was much delayed, with the bulk of the activity occurring during the rinse; DRZ had no effect on this response. **B**. Superimposed bar graphs show average responses (±SEM) across all B6 neurons (N=11) before (blue) and after (red) DRZ treatment. ANOVA yielded a significant effect of stimulus (*p* < 0.0001) but no effect of DRZ (*p* = 0.66) or a stimulus X DRZ interaction (*p =* 0.36). **b1-b3**. Responses of each neuron across different clusters (N_as_, EG_Nai_ /EG_acid_, and sugar). Despite the lack of the effect with an overall ANOVA, responses to salt, salt plus amiloride, and all sugars were consistently suppressed in EG_Nai_ /EG_acid_ cluster neurons but not in the other (N_as_ and sugar) cluster types. **C**. Superimposed bar graphs show average responses (±SEM) across all KO1+3 neurons (N=7) before (blue) and after (red) DRZ treatment. ANOVA yielded a significant effect of stimulus (*p* = 0.012) and an interaction between stimulus X DRZ (*p =* 0.002). **c1 and c2**. Effect of DRZ in the N_as_ and EG_Nai_ cluster neurons of KO1+3 mice. **D**. Bar graph (means±SEM) representing the average responses to all the stimuli across different clusters after combining B6 and KO1+3 mice (ANOVA: DRZ-*p=*0.141, cluster and stimulus= *p’s* < .0001, DRZ X cluster, DRZ X stimulus, DRZ X cluster X stimulus – *p’s* < .0001). Interestingly, DRZ suppressed salt and sugar mediated responses of EG_Nai_ cluster neurons. Post-hoc paired Bonferroni-adjusted t-tests showed significant reductions for N100, N100+a, and each sugar (all *p’s* < 0.05) in the EG_Nai_ group but not for bitter (B3) or acid (I10) responses.

**Figure 5B** illustrates the effect of DRZ on mean taste responses in B6 mice (N=10, left panel). Across all stimuli, there was not a significant suppression by DRZ, nor was there an interaction between stimulus and DRZ (see caption for all DRZ statistics). However, we noted that the small amiloride-insensitive sodium responses seemed obliterated by DRZ treatment, and that there was a trend for reduction in the sugar responses. Thus, we suspected that neuron-type specific effects were obscured. Because there were not enough neurons in this sample to perform statistical analyses as a function of chemosensitive type, individual responses were examined separately for the different types (b1-b3). Using this approach, it was clear that DRZ consistently reduced NaCl and sugar responses, but only in the electrolyte generalist, amiloride-insensitive groups (EG_Nai_ and EGacid). Bitter and acid responses did not change systematically, regardless of neuron group. **Figure 5C** illustrates the DRZ effect in KO1+3 mice. In contrast to B6 mice, across all neurons (N=8, left panel), the stimulus X DRZ interaction was significant. Moreover, when segregated by cluster, effects were identical to those in B6 mice: sugar and NaCl responses were consistently reduced in the EG_Nai_ / EGacid groups but not the Nas group (there were no KO1+3 sugar-sensitive cells). To garner enough neurons to statistically compare neuron types, we combined WT and KO1+3 cells (**Fig. 5D**). This analysis yielded main effects of DRZ, cluster, and stimulus, as well as interactions between these variables. Post-hoc comparisons show significant reductions for N100, N100+a, and each sugar in the EG_Nai_ group, but not for bitter (B3) or acid (I10) responses. Moreover, there was no effect on responses to any stimulus in the other clusters, including those to sugars in the sugar cluster. This finding suggests that one mechanism for T1R-independent responses elicited by hyperosmotic concentrations of sugars is the result of activating a shared carbonic anhydrase-mediated mechanism in amiloride-insensitive type III taste bud cells.

### T1R1+3-heterodimer contributes to umami synergism, but an antagonist for mGluR4 does not affect residual glutamate responses

Because the T1R1 + T1R3 heterodimer is the major taste receptor for amino acids, we also evaluated glutamate responses in a subset of neurons. In our previous NST study, similar to the case for sugars, genetic deletion of T1R1+T1R3 failed to eliminate glutamate responses. Although KO1+3 mice displayed significantly attenuated responses to 100 mM MSGa + IMP (umami), responses to 600 mM umami or MSGa alone were unaffected (Kalyanasundar et al., 2020). Here, we used additional concentrations (50, 300, and 600 mM) of umami and MSGa to gain a fuller picture of T1R-independent glutamate responses. **Figure 6A** shows that activity elicited by MSGa increased with stimulus concentration but did not vary by genotype (ANOVA: strain, *p* = 0.935, concentration, *p* < 0.00005, strain X concentration, *p=*0.076). In contrast, for umami, there was a major effect of genotype (ANOVA: strain, *p* = 0.008, concentration, *p* = < 0.00001, strain X concentration, *p =* 0.0035). Post-hoc tests indicated that 50 (*p =* 0.0002) and 300 (*p =* 0.007) mM umami responses were smaller in the KO’s but 600 mM responses were comparable (*p =* 0.12). IMP by itself (presented in a subset of cells) elicited small responses even in B6 mice and these were negligible in the KO’s (*p* = 0.02). We also quantified the synergistic ratio ([umami response/sum of responses to MSGa+IMP], (Tokita et al., 2012), **Fig. 6B**). The average synergistic ratio exceeded the standard criterion (1.2) for the two lower concentrations but only for B6 mice. At 600 mM, synergy was not apparent for either strain (ANOVA: strain: *p =* 0.01, concentration: *p =* 0.63, and concentration X strain: *p* = 0.06). To evaluate which cell types carried the T1R1+3-independent glutamate information, responses were plotted separately by cluster type and strain (**Fig. 6D**), using the cluster groups defined in **Figure 2**. There were responses to umami and MSGa in each cluster, but enhancement of the MSGa response by IMP was only apparent in the sugar cluster; here called “sugar/umami” because they responded robustly to both sugars and MSGa+IMP (pink circles, **Fig. 6D left panel**). As noted above, this cluster was present only in WT mice, suggesting that these umami responses are T1R-dependent. However, glutamate and umami not only stimulated sugar/umami cluster neurons but also robustly activated EGacid and EG_Nai_ neurons, particularly at concentrations above 50 mM. In these clusters, glutamate responses were comparable for the two strains (ANOVAs; clusters: *p* = <0.0001, stimulus; *p* < 0.0001, strain: *p =* 0.37, stimulus X strain: *p =* 0.07, cluster X strain: *p =* 0.77, and stimulus X cluster X strain: *p* = 0.69), suggesting a T1R-independent basis.

**Figure 6.**
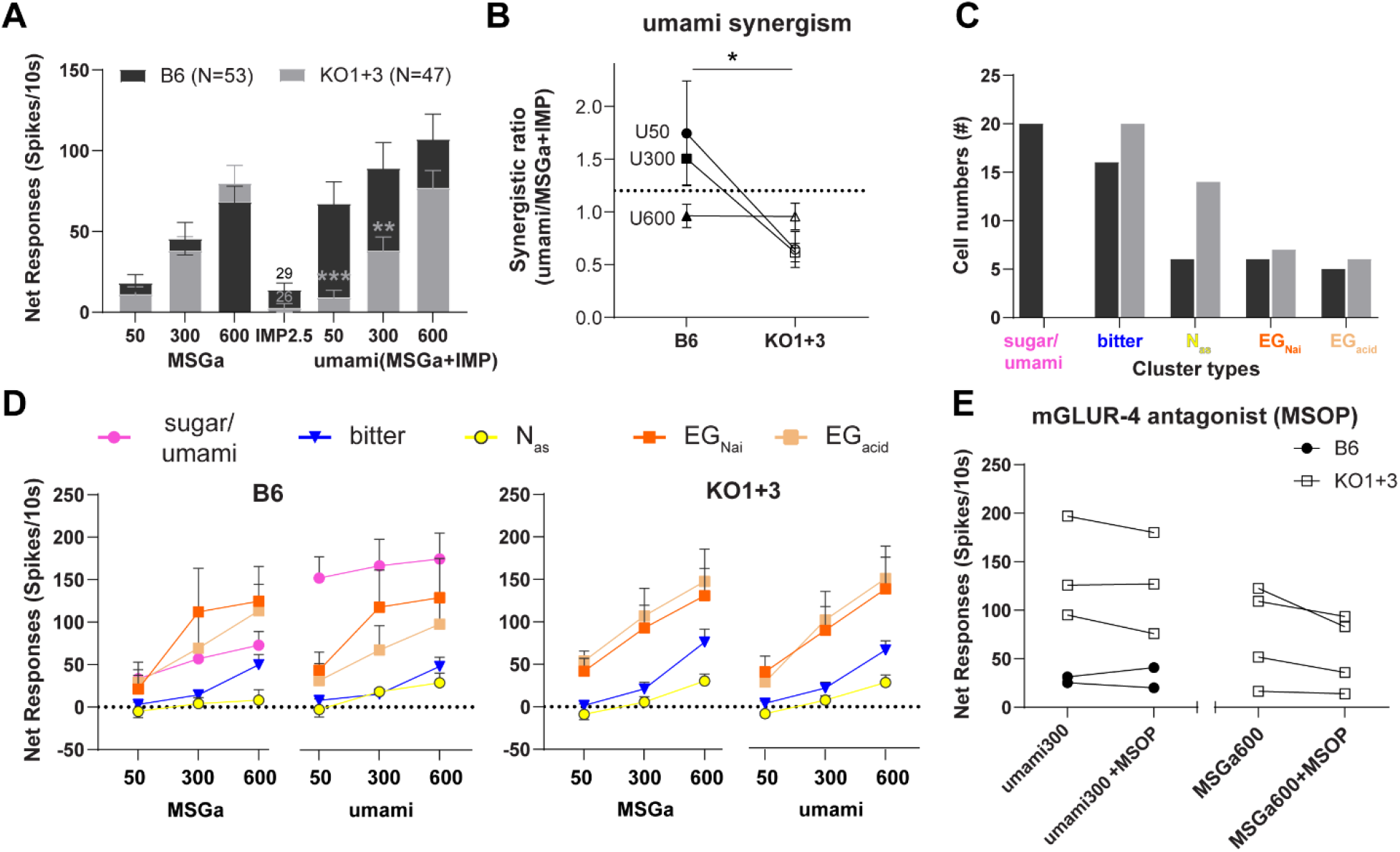
Enhancing neuronal response to glutamate by IMP requires the T1R1+T1R3 heterodimer, but an mGluR4 antagonist did not affect T1R-independent glutamate responses. **A**. Superimposed bar graphs represent the mean (±SEM) responses of PBN neurons in B6 (dark bars) and KO1+3 (grey bars) mice elicited by monosodium glutamate with amiloride (MSGa) and, inosine 5’-monophosphate (IMP), and the MSGa + IMP mixture (umami). ** & ***-P<0.001 and < 0.0001 post-hoc between strains. **B**. Synergistic ratio (means±SEM) across three different concentrations of umami compared between B6 and KO1+3 mice. Only cells tested with IMP were included (N’s: B6 vs. KO1+3, 29 vs. 26). The horizontal dotted line at 1.2 on the y-axis is the previously (Ninomiya and Funakoshi, 1989; Tokita et al., 2012) established criterion for umami synergism. Umami synergism at 50 and 300 mM was largely reduced in KO1+3 mice compared to B6 (ANOVA: effect of strain, *p* = 0.012). Note that response to umami at 600 mM did not show any increased responsiveness to IMP addition. **C**. Cluster classification of PBN neurons tested with all the concentrations of umami and glutamate across B6 (black bars) and KO1+3 (grey bars) mice. Clusters are based on the analysis in **Figure 2**. The sugar/umami neurons are the same as those called “sugar” in the preceding Figures. Note KO1+3 mice lacked sugar/umami best neurons. **C**. Line graph representing average net responses (± SEM; for clarity, only top error bars are shown) to the three concentrations of MSGa and umami across five cluster types for B6 (left) and KO1+3 mice (right). Umami synergism was evident in B6 mice sugar/umami neurons (pink filled circles) at all concentrations (Bonferroni adjusted paired t-tests, MSGa vs umami at all concentrations, *p’s* < 0.0001). Notably, there appeared to be a ceiling effect for the umami response at the lowest concentration tested (50 mM). In contrast, in EG_Nai_ and EG_acid_ cluster neurons (both strains) umami and MSGa responses were comparable. **D**. The contribution of the metabotropic receptor (mGluR) for the T1R-independent glutamate-mediated neuronal responses was tested using a selective antagonist of mGluR4 (MSOP at 0.5 mM; (Pal Choudhuri et al., 2016) by mixing with umami (300 mM) and glutamate (600 mM). Though only a small sample of cells was tested (N’s: KO1+3: 3-4 and B6: 2), the mGluR antagonist only produced a small, non-significant suppression.

Previous studies suggest that metabotropic glutamate receptors (mGluRs) may be responsible for the T1R-independent glutamate responsiveness (Chaudhari et al., 1996; Chaudhari and Roper, 1998; Chaudhari et al., 2000). To test this hypothesis, we used an mGluR4 antagonist, MSOP (α-Methylserine-O-phosphate), as a cocktail with umami (300 mM) or glutamate (600 mM), depending upon which response was larger in a particular cell (**Fig. 6E**). However, the addition of MSOP did not suppress these responses. Thus, our results suggest that some T1R-independent amino acid responses are also mGluR-independent.

### B6 and KO 1+3 neurons with similar anatomical locations and receptive fields were sampled

Figure 7. shows the distribution of PBN recording sites for each of the two strains of mice. Neurons were sampled primarily in the “waist region”; i.e., the ventral lateral and central medial subnuclei. Their distribution appeared to correspond with that for SATb2-labeled neurons. Cells spanned an anterior-posterior region extending from the caudal pole of the nucleus ~400 μM rostrally to the level where the external lateral nucleus begins to form, with the greatest proportion of neurons identified at the caudal level (49%), and progressively fewer at the middle (33%) and most rostral (18%) levels within this region. Comparable proportions of B6 and KO 1+3 neurons were encountered across these three levels (*p* = 0.648, χ2). There was no obvious chemotopic organization. Neurons with receptive fields (RF’s), classified as anterior (A) only, posterior (P) only, or mixed (M), of the B6 and KO 1+3 cells also were equivalent for the two strains (*p* = 0.198, χ2). Across both strains, a majority of neurons (65%) had RFs confined to the A (anterior tongue and/or nasoincisor ducts), and 13% had RFs confined to the P (soft palate, foliate papillae, circumvallate papilla) or a combination from the A and P (22%). There was a non-random relationship between chemosensitivity and receptive field (*p* = 0.003, χ2); most strikingly, 80% of Nas neurons had A RFs and none had P RFs whereas bitter cells had RFs more evenly split between A only (39%), P (32%), and mixed (29%).

## Discussion

It has been known for two decades that the T1R receptor family forms heterodimers expressed in taste bud cells that are the principal receptors for responsiveness to umami-like stimuli (T1R1+T1R3) and sweeteners (T1R2+T1R3). Yet, residual sensitivity to T1R ligands behaviorally (Damak et al., 2003; Zhao et al., 2003; Delay et al., 2006; Treesukosol et al., 2009; Treesukosol et al., 2011b, a; Treesukosol and Spector, 2012; Smith and Spector, 2014; Blonde and Spector, 2017; Blonde et al., 2018; Schier et al., 2019; Yasumatsu et al., 2020) physiologically (Smith and Spector, 2014; Glendinning et al., 2015; Yasumatsu et al., 2020), and neurally at the level of the peripheral gustatory system (Damak et al., 2003; Zhao et al., 2003; Lemon and Margolskee, 2009; Yasumatsu et al., 2020) and NST (Lemon and Margolskee, 2009; Kalyanasundar et al., 2020) has been documented in mice lacking one or both heterodimers. The present study extends previous observations of T1R-independent sugar and glutamate responses to the PBN, provides further insights into their functional organization, and clarifies what taste receptor mechanisms are and are not contributing. Similar to NST, there was a notable absence of sugar/umami selective neurons in the KOs. Nevertheless, both classes of stimuli evoked salient neural activity. Indeed, sugar-elicited T1R-independent responses were dramatically larger than in the medulla. In KO mice, hyperosmotic sugars primarily activated cells broadly tuned to electrolytes; nearly identical responses occurred in electrolyte-generalist WT neurons. Like Type III taste bud cells and their recipient CT neurons, a carbonic anhydrase inhibitor, dorzolamide (DRZ), suppressed NaCl responses in EGacid and EG_Nai_ PBN cells (Oka et al., 2013; Lewandowski et al., 2016). More surprisingly, DRZ suppressed sugar responses in these neurons, suggesting a novel transduction mechanism for T1R-independent responses to hyperosmotic sugars originating in Type III taste bud cells. In contrast, the sensory receptors contributing to residual umami responsiveness in KO mice remains unclear.

**Figure 7.**
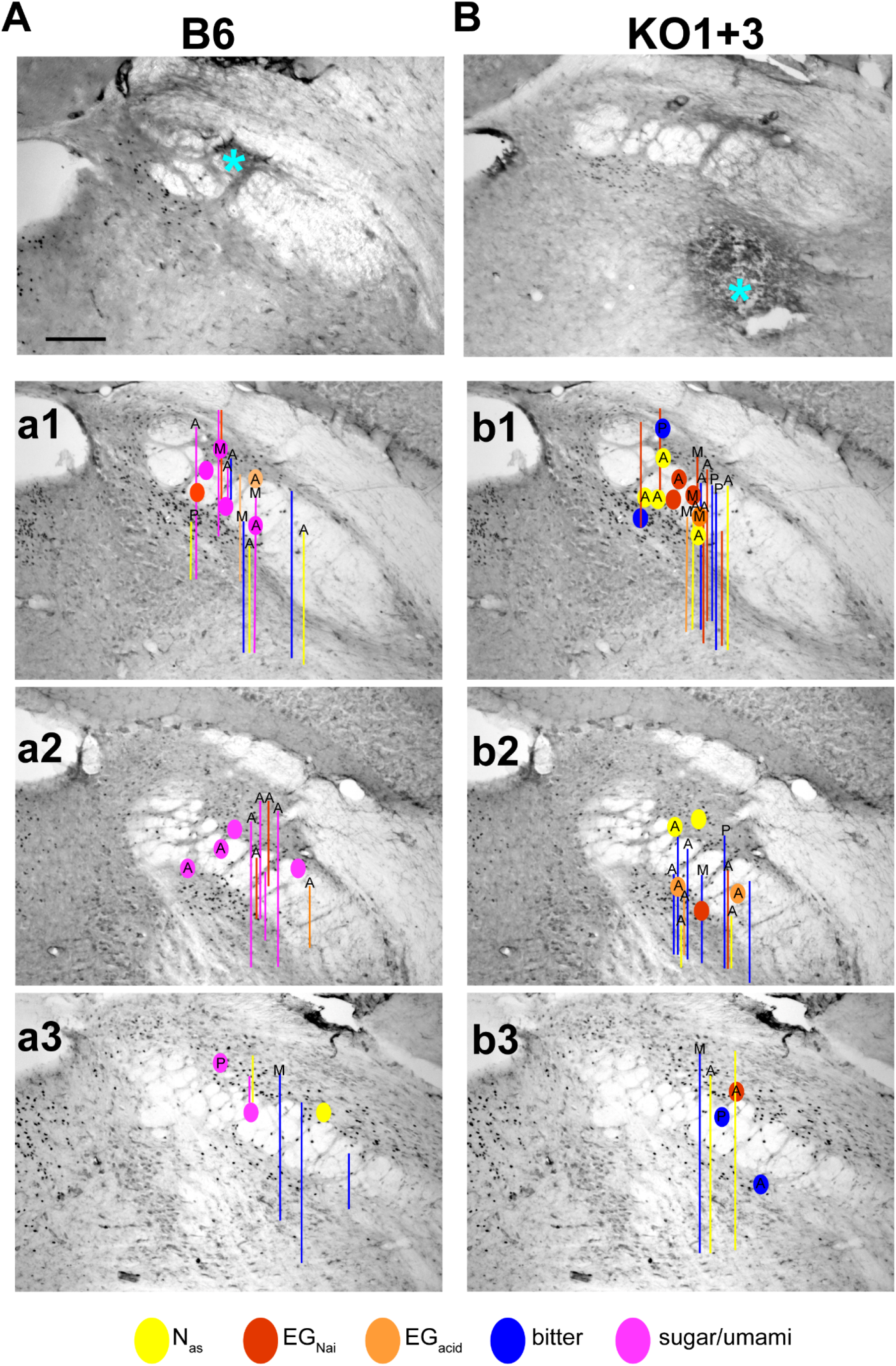
Histological analysis of recording sites on Satb2 stained sections of taste PBNA. Brightfield photomicrograph of a Satb2-immunostained section depicts an electrolytic lesion (asterisks) made at the site of recording where a sugar cluster neuron was recorded from a B6 mouse. Satb2+ neurons are evident by the dark nuclear staining. Lesions in SAT2b DAB-immunostained tissue were apparent because they produced dark nonspecific staining, presumably due to the anti-mouse secondary antibody for Satb2. Scale bar, 100 μm. **a1-3**. Reconstruction of recording locations in B6 mice at three different representative (caudal to rostral) levels of taste PBN superimposed on a series of PBN sections immunostained with Satb2. Symbols represent lesions made at the site of recording and lines represents the estimated trajectory of tracks through the nucleus when lesions were made dorsal or ventral to the recording site. The top of the line is the estimated site of recording. Symbols and lines are color-coded based on chemosensitive neuron clusters. The letters inside the symbols or at the top of the tracks denote receptive fields [anterior (a), posterior (p), or mixed (m)]. **B**. An example of a Satb2-immunostained section from a KO1+3 mouse where we made a lesion (asterisk) ventral (400μ) to the sites of recording for two different cells: a bitter neuron and an EG_Nai_ neuron. The lesion was made beneath the PBN in a location where responses to depressing the mandible were evident. **b1-b3:** KO1+3 mice recording sites and trajectories are symbolized in the representative sections as for B6 mice. It is worth noting that the distribution of the PBN neurons we recorded from roughly correspond that for the Satb2 labelling.

### Central representation of T1R-independent glutamate responses

The most notable effect of T1R1+T1R3 deletion on glutamate responses was the lack of conspicuous enhancement of the MSGa response by IMP, consistent with previous observations in the CT and NST (Damak et al., 2006; Lemon and Margolskee, 2009; Kalyanasundar et al., 2020). In the NST of WT mice, IMP enhanced MSGa responses dramatically (320%) but there was also a small augmentation (20%) of MSGa responses by IMP in the T1R KO’s (Kalyanasundar et al., 2020), of interest since KO 1+3 mice can discriminate MSGa + IMP, but not MSGa, from water (Smith and Spector, 2014; Blonde et al., 2018). In the pons, there was likewise a large IMP enhancement of MSGa responses in the WT’s. However, there was none in T1R KO’s, suggesting that synaptic processing may filter out the small increase in the medulla. Thus, an explanation for the efficacy of IMP in facilitating behavioral detection of MSGa remains elusive. T1R1 + T1R3 knockout reduced MSGa + IMP responses but had no impact on MSGa responses. Earlier work suggests that mGluR receptors contribute to T1R-independent glutamate responses (Chaudhari et al., 1996; Chaudhari and Roper, 1998; Yasumatsu et al., 2012; Kusuhara et al., 2013). In the current study, treatment with the mGluR4 antagonist, MSOP, only produced a non-significant trend for a decrement in glutamate responses. Because the sample size was small and we did not test multiple antagonist concentrations, nor an mGluR1 antagonist, these results are not definitive. On the other hand, aside from umami responses in WT sugar/umami neurons, MSGa + IMP and MSGa mainly stimulated EG_Nai_ and EGacid cells, consistent with the conclusion that glutamate is not the constituent of the MSG molecule responsible for eliciting the observed T1R-independent responses.

### Central representation of T1R-independent sugar responses

Previous neurophysiological studies in T1R3 KO mice report prominent reductions in responses to monosaccharides, disaccharides, and artificial sweeteners, but some persistent sugar responsiveness (Damak et al., 2003; Zhao et al., 2003; Lemon and Margolskee, 2009; Yasumatsu et al., 2020). The present findings are broadly consistent with these observations. However, in the CT, glucose responses appear less susceptible to T1R3-KO than those to sucrose (Damak et al., 2003; Yasumatsu et al., 2020) or fructose (Damak et al., 2003). In our NST study it similarly appeared that T1R-independent glucose (though also fructose) responses were more salient (Kalyanasundar et al., 2020). However, we used the monosaccharides at a higher (1000 mM) concentration than sucrose (300 mM). Here, T1R-independent sucrose responses were just as prominent when we tested sugars at an equimolar (600 mM) concentration. Similarly, Lemon observed no difference between T1R3-independent responses to 1000 mM glucose and sucrose in rNST (Lemon and Margolskee, 2009). Another contrast in peripheral vs central sugar responsiveness is that selective “sugar-best” CT neurons persist in KOs, though fewer in number and less responsive than in WT animals (Yasumatsu et al., 2012; Yasumatsu et al., 2020). Such neurons were not observed in the NST (Lemon and Margolskee, 2009; Kalyanasundar et al., 2020) or PBN.

### Potential alternative sugar sensing mechanisms

Prior work suggests that glucose transporters comprise part of an alternative sugar-sensing mechanism (Yee et al., 2011) that can also detect glucose-containing disaccharides since α-glucosidases in taste bud cells digest disaccharides (Sukumaran et al., 2016). Previous studies demonstrated that a blocker for the sodium-glucose transporter (SGLT), phlorizin, attenuates sucrose and glucose CT nerve responses and raises the glucose detection threshold in human subjects (Yasumatsu et al., 2020; Breslin et al., 2021), suggesting a role for this transporter. However, phlorizin had no effect on sugar-elicited PBN responses. The reason for this disparity is unclear. Although an effect may have emerged with more neurons, the carbonic anhydrase inhibitor, dorzolamide (DRZ), produced a robust decrease in a similar size sample. Notably DRZ only affected sugar responses in amiloride-insensitive electrolyte generalist neurons, not sugar/umami cells. DRZ also affected NaCl, but not acid responses in these neurons. These effects on NaCl parallel those in Type III taste bud cells and presumed recipient CT fibers (Oka et al., 2013; Lewandowski et al., 2016), implying that the DRZ-suppressible PBN sugar responses arise from the same taste bud cell type and share components of the transduction cascade for amiloride-insensitive NaCl responses. The transduction mechanism(s) for amiloride-insensitive NaCl responses is poorly understood but existing observations provide some insight (Spector and Travers, 2021). Notably, sugar responses in EG_Nai_ and EGacid cells occurred at concentrations of 600 mM and higher but were small or absent at 300 mM (**Fig. 3**), suggesting that a hyperosmotic concentration was critical. Interestingly, Lewandowski and colleagues showed that hyperosmotic conditions magnified amiloride-insensitive NaCl responses in a subpopulation of Type III cells. The carbonic anhydrase enzyme family participates in pH regulation, ion and fluid transport by catalyzing the reversible hydration of CO2 to yield H+ and HCO3-(Sterling and Casey, 2002; Supuran, 2021). In Type III taste bud cells, carbonic anhydrase IV, attached to the extracellular membrane, accounts for the “taste” of carbonation by generating protons from CO2 that pass through the OTOP channel ultimately depolarizing the cell (Chandrashekar et al., 2009; Lossow et al., 2017; Tu et al., 2018). One possibility is that hyperosmotic sugars generate a similar CA-dependent pH or ionic shift that then activates Type III cells.

### Functional significance and perspectives

Here we show robust T1R-independent responses in the PBN, an essential relay in the ascending rodent gustatory pathway (Norgren and Leonard, 1971, 1973), elicited by hyperosmotic sugars. It seems doubtful that these responses, that occur in electrolyte-generalist neurons in both KO and WT mice, relate to “sweetness” or the hedonic value of sugars. The order of effectiveness of different sugars in activating these cells corresponded poorly to their efficacy in eliciting licking, whereas activity in selective sugar/umami WT neurons corresponded well (**Fig. 3**). Indeed, T1R KO mice exhibit severe deficits in sugar discrimination and display negligible concentration-driven licking to these compounds in the absence of post-oral reward (Zhao et al., 2003; Zukerman et al., 2009; Treesukosol et al., 2011b, a; Treesukosol and Spector, 2012; Smith and Spector, 2014). Based on broad sensitivity across sugars, it is also unlikely these responses are a substrate for the behavioral ability of mice learn a glucose versus fructose discrimination (Schier and Spector, 2016; Schier et al., 2019) and improbable that they account for the special status of glucose in eliciting cephalic phase insulin release (Grill et al., 1984; Glendinning et al., 2018). In fact, because a hyperosmotic sugar concentration appeared necessary, we suspect other hyperosmotic stimuli would be efficacious. Nevertheless, while these responses are unlikely to arise from a sugar-specific receptor, hyperosmotic sugars are physiological stimuli, routinely included in behavioral taste tests, and encountered in the human diet. It was striking that these tastants elicited responses that rivaled those to 100 mM NaCl in the EG_Nai_ electrolyte generalist cells; in fact, they were often the nominal “best-stimulus” (**Fig. 2**). Thus, though not a sugar-specific signal, it seems feasible that these potent responses could comprise cues that T1R double-KO mice could utilize, for example, to associate with post-oral reward to support preference (Treesukosol et al., 2009; Treesukosol et al., 2011a; Zukerman et al., 2011; Sclafani, 2013; Tan et al., 2020).

That hyperosmotic PBN sugar responses likely originate in Type III taste bud cells is provocative in light of the discovery of Type III “BR” (broad) acid-responsive cells that also have intrinsic sensitivity to sweet, umami, and bitter stimuli. Inhibiting BR responses impairs hedonic responses to taste stimuli (Dutta Banik et al., 2020). Another recent study shows that yet a different mechanism for detecting glucose, glucokinase, is expressed primarily in Type III cells (Chometton et al., 2022). Together these peripheral and central data emphasize the multisensitive nature of certain gustatory cells and the complexity of sugar-detecting mechanisms.

## Acknowledgments

This work was supported by **NIH DC004574 to A.C.S**. Thanks to Dr. Joseph B. Travers for his valuable comments on the initial manuscript. The excellent technical assistance of Cemaliye Semmedi, Andrew Harley, and Charlotte Klimovich are greatly appreciated. We thank the Zuker laboratory for providing the original breeding pairs to the Spector laboratory for generating the dbl KO’s.

